# Foldy: a web application for interactive protein structure analysis

**DOI:** 10.1101/2023.05.11.540333

**Authors:** Jacob B. Roberts, Alberto A. Nava, Allison N. Pearson, Matthew R. Incha, Luis E. Valencia, Melody Ma, Abhay Rao, Jay D. Keasling

**Author notes:** Co-first authors (Jacob B. Roberts, Alberto A. Nava).

## Abstract

Foldy is a cloud-based application that allows non-expert scientists to easily access and utilize advanced AI-based structural biology tools, including AlphaFold and DiffDock. Built on Kubernetes, it can be deployed by universities, departments, and labs without requiring hardware resources, but can also be configured to utilize available computers. Foldy enables scientists to predict the structure of proteins and complexes up to 3000 amino acids, visualize Pfam annotations, and dock ligands with AutoDock Vina and DiffDock.

Our manuscript describes the user interface and deployment considerations of Foldy, as well as some of our applications. By democratizing access to sophisticated AI-based tools, Foldy can facilitate life science research and promote the wider adoption of structural bioinformatics tools. Our work demonstrates that even the most advanced tools can be made accessible to a broad audience through user-friendly platforms like Foldy, and we believe it will be a valuable resource for researchers across scientific disciplines. The public structures available on the Lawrence Berkeley Labs Foldy deployment can be viewed at https://foldy.lbl.gov.

**Author Summary:** Foldy is a cloud-based application that enables scientists to use AI-based structural biology tools such as AlphaFold and DiffDock without software expertise. Built on Kubernetes, it can be set up by universities, departments, and labs with no need for hardware resources. Foldy can predict the structure of proteins and complexes up to 3000 amino acids, visualize Pfam annotations, and dock ligands with AutoDock Vina and DiffDock. Our public structures can be viewed at https://foldy.lbl.gov.

Our manuscript highlights the user interface, deployment considerations, and product applications of Foldy. It’s an accessible solution for researchers who are not software experts and can handle the traffic of thousands of users and hundreds of thousands of protein structures and docked ligands. This makes advanced AI-based tools more widely available, paving the way for accelerating life science research.

By developing an easy-to-use platform, our work demonstrates that even the most sophisticated AI-based tools can be made accessible to a wide audience. Foldy enables more scientists to draw from the rapidly growing field of structural biology, making it a valuable tool for researchers across scientific disciplines. We look forward to its adoption by the scientific community.

## Introduction

Recent advances in machine learning have led to the development of highly accurate protein structure prediction methods [1–4], but their adoption has largely been limited to computational biologists. These methods have produced impressive results in numerous applications including *de novo* protein design [5] and protein-protein interaction screening [6]. However, the steep requirements for storage space, GPU processing power, and RAM make the use of these tools difficult for many end users. Nvidia created the BioNeMo service to make AI more accessible to life science researchers, but it is a private implementation and currently only available to a few biotechnology companies. Programs such as ColabFold [7] and AlphaFold-Colab [8] have been developed to meet this need by providing custom Google Colaboratory Jupyter notebooks which utilize free compute resources hosted by Google Cloud. These Jupyter notebooks provide an interactive mode of using AlphaFold without the need for any complex installation or configuration. However, there are several limitations to these notebooks including session timeouts, limited GPU power, and limited batch processing capabilities. These issues are exacerbated when large proteins (>1000 amino acids) are modeled which have higher resource demands. Ultimately, these limitations mean that scaling up to more than a handful of structures is prohibitively difficult.

Here we present Foldy, an easy-to-deploy and easy-to-use modern web app for folding a protein (AlphaFold[1]), predicting domain annotations (Pfam[9]), and docking small molecule ligands (AutoDock Vina[10] or DiffDock[11]). Its primary objectives are to provide an intuitive interface, facilitate deployment for IT administrators, and enable prediction tasks for tens to thousands of users per instance. The integrated tools within Foldy facilitate a seamless transition between protein structure prediction and downstream analysis. It can be rapidly deployed in a cloud environment, and also offers the possibility of a hybrid deployment utilizing local computational resources. The design of the Foldy architecture aims to lower barriers to entry for both end users and institutions.

## Results

### Interface

Foldy has four main views: the New Structure view, the Dashboard, the Tag view, and the Structure view. The New Structure view (Fig S1) is where users can submit new structure prediction tasks. At a minimum, users must provide an amino acid sequence and a name for the structure. The Dashboard (Fig 1) serves as the app’s landing page, providing access to all other pages. By default, the Dashboard displays a table of the user’s structures, but a search bar allows users to filter structures by name, user, protein sequence, or tag. The Tag view (Fig S2) displays all structures with a particular tag, and exposes bulk tasks such as downloading structures or docking small molecule ligands.

**Fig 1:**
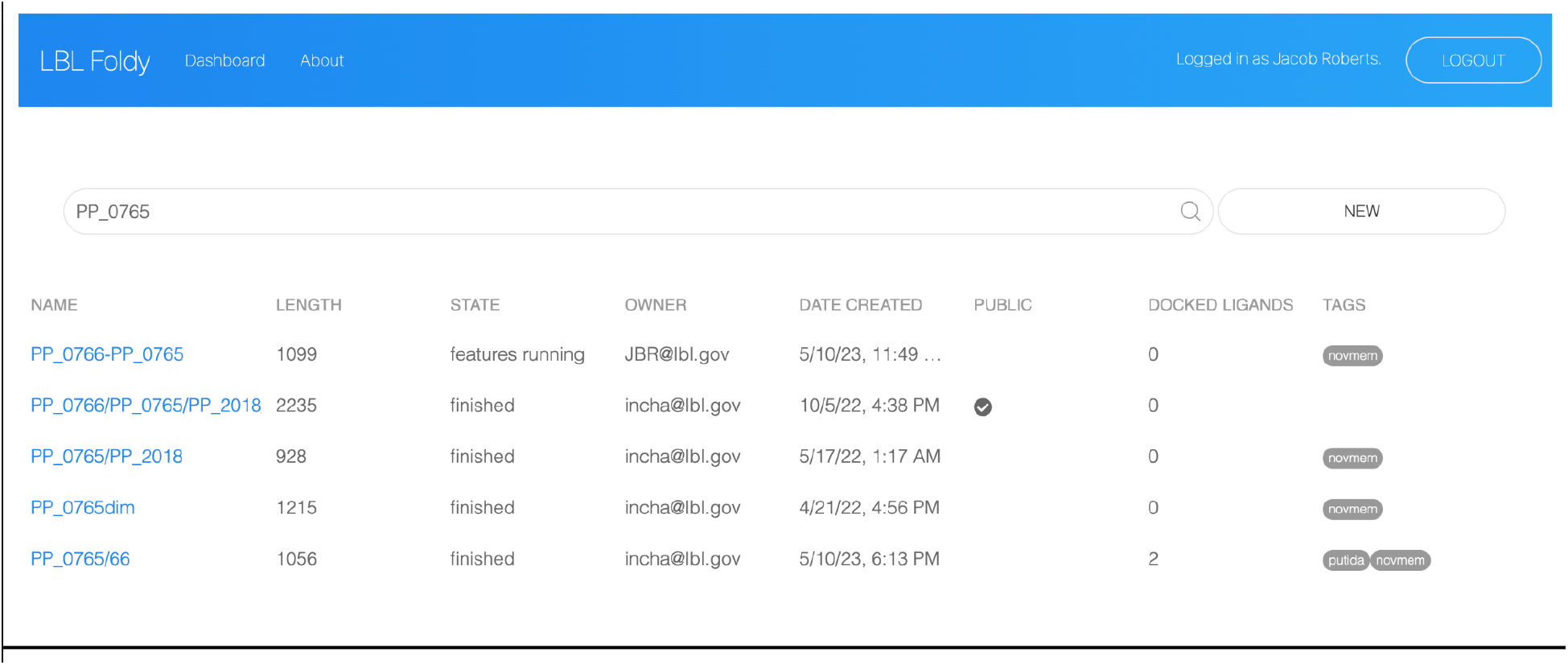
Dashboard View. This is the landing page after authentication displaying a table of fold tasks. The table includes information about the status of each fold. The search bar at the top of the page enables searching and filtering folds based on their name, sequence, or tag.

There are two types of users: editors and viewers. Editors have full read and write access. Viewers are any user with a Google account, and are allowed view-only access to structures which have been explicitly marked “public” and their associated data (logs, docking runs). Users are authenticated by their Gmail account, and user types are flag controlled. You can view our public structures at https://foldy.lbl.gov.

The Structure view has two columns: the predicted structure is on the left and a tool panel is on the right (Fig 2). By default the tool panel displays the amino acid sequence (Fig 3A), and Pfam domain annotations can be overlaid on both the structure and sequence(s) (Fig 2 left, Fig 3A). A number of actions are available to users through the tabs in the tool panel. For example, users can predict residue interactions and complex formation using contact probability maps (Fig 3E) [6]. Users can segment proteins into domains and predict inter-domain flexibility using the Predicted Alignment Error (PAE, Fig 3D) [3]. Additionally, users can dock small molecule ligands with AutoDock Vina or DiffDock by specifying the SMILES string and optionally a bounding box around a residue (Fig 3F) [10,11].

**Fig 2:**
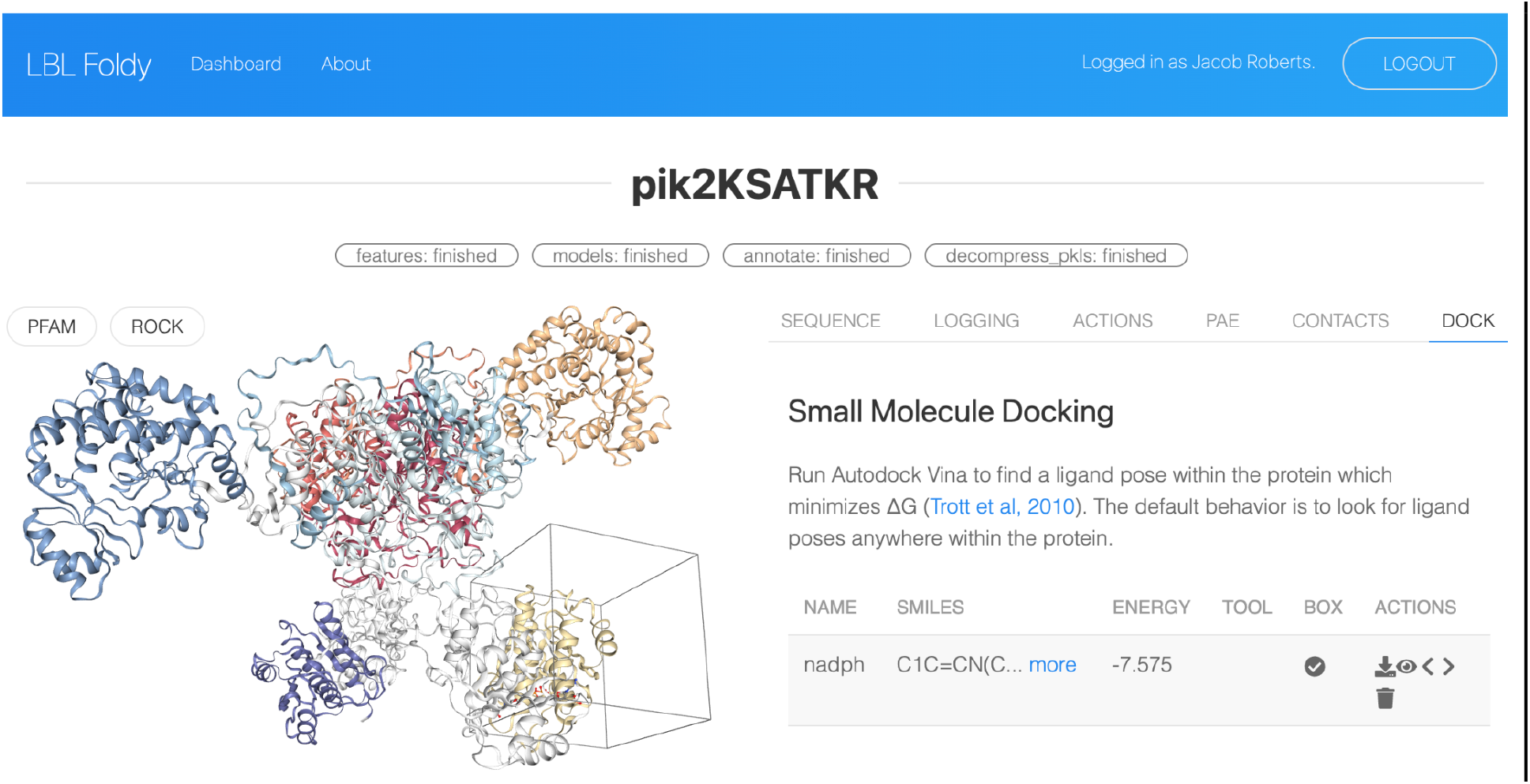
Structure View. The Structure view displays an example predicted structure of a homo-dimeric polyketide synthase module with the docking tool selected. Pfam domains are highlighted on the peptide structure. We docked NADPH with the docking tool, and its predicted binding affinity and pose are displayed.

**Fig 3:**
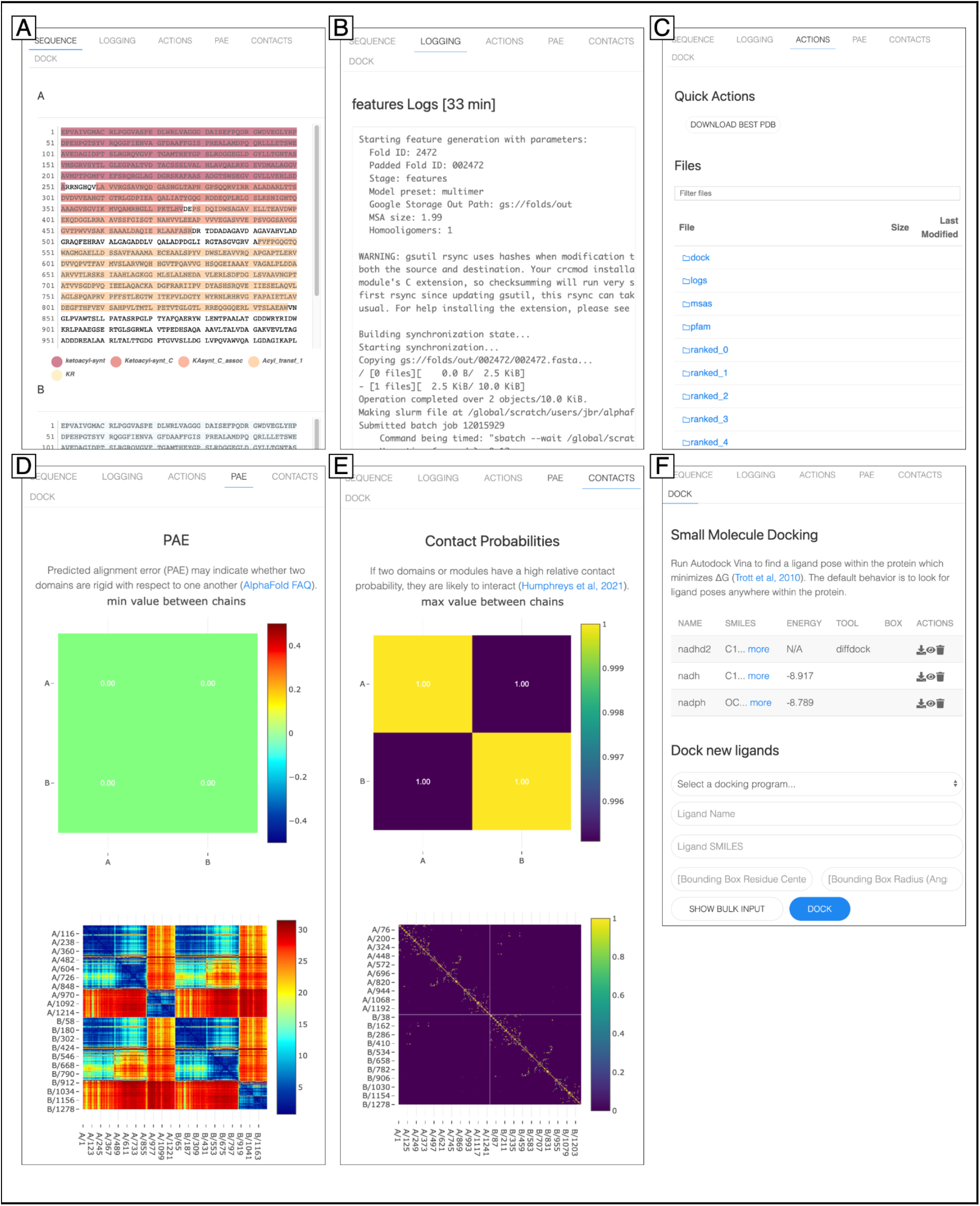
Structure View Toolbar Tabs. ***A. Sequence***: this tab displays the peptide sequences, optionally annotated with domain predictions, such as the pfam annotations displayed here. ***B. Logging***: this tab displays real-time logs of each sub-task within a protein structure prediction task. ***C. Actions***: this tab allows a user to download the top ranked pdb structure file of a given fold as well as all the intermediate files generated by the fold pipeline ***D. PAE***: this tab shows the predicted alignment error heatmap, measured in Ångstroms, between each residue and between each chain. The PAE is derived from the AlphaFold model. ***E. Contact Probability***: this tab shows the contact probability heatmap between each residue and between each chain. The contact probabilities are derived from the AlphaFold model. ***F. Docking***: this tab shows a table with any docking results as well as a submission form for new docking tasks. The docking submission form requires a ligand name and SMILES string, and optionally a bounding box can be specified of a particular radius around a particular residue (Vina only).

### Facile Deployment

A local version of Foldy can be set up in seconds with Docker Compose, but none of the analytical tools are supported locally. The full Foldy can be deployed in the cloud in a matter of hours by an IT administrator. It requires a few manually provisioned resources: a web domain for the app, a static IP address, a database, and a Kubernetes cluster. The remainder of the app, including the frontend, backend, and compute workers, are all managed by a single Helm chart. Helm is a command line tool for managing Kubernetes configuration, and with one command can deploy all the necessary Kubernetes resources. The infrastructure schematic is illustrated in Figure S3, and the Helm chart with corresponding step-by-step deployment instructions are available at https://github.com/JBEI/foldy.

### Scalability

By default, Foldy uses cloud compute for all tasks, meaning any lab or institution, regardless of their hardware, is able to set up Foldy. In this setup, jobs are run on ephemeral machines in the cloud that are spawned when work tasks are queued and deleted when the work is complete. The work tasks are tracked in a queue (implemented with RedisQueue), and the worker machines are automatically created and destroyed by Kubernetes (implemented with Prometheus and KEDA). Importantly, each worker machine can be provisioned with high-memory GPUs, enabling the prediction of large protein structures.

Institutions with access to their own compute resources, including compute clusters, can run the worker threads on their own machines. To run a worker on a local compute resource which supports docker, one can run the “worker” docker image on the machines with the appropriate flags, and set up tunnels to the cloud databases. To use local compute resources which don’t support docker, one can create bash scripts which execute each tool. For example, to run AlphaFold jobs on a university cluster which does not support docker, one must create a variant of “worker/run_alphafold.sh” which invokes the local AlphaFold installation.

## Discussion

This tool greatly facilitates research because it makes complex tools accessible. The LBL Foldy instance (https://foldy.lbl.gov) has been used by 55 researchers across 6 labs, to predict 4802 structures and dock 2754 ligands. It has been used in over a dozen projects, three of which are case-studied below.

### Foldy Case Studies

One researcher was interested in changing the substrate preference of a long-chain fatty acyl-AMP ligase (FAAL) from long-chain to medium-chain. They used Foldy to predict the wild type protein’s structure, and used AutoDock Vina to dock octanoyl-AMP in the active site tunnel. They considered two point mutations to tryptophan which they hypothesized would obstruct the substrate tunnel and shorten chain-length preference of the FAAL. They found from the structure that one tryptophan point mutation would occlude even C8 compounds, while the other would occlude C10 or larger (Fig 4).

**Fig 4:**
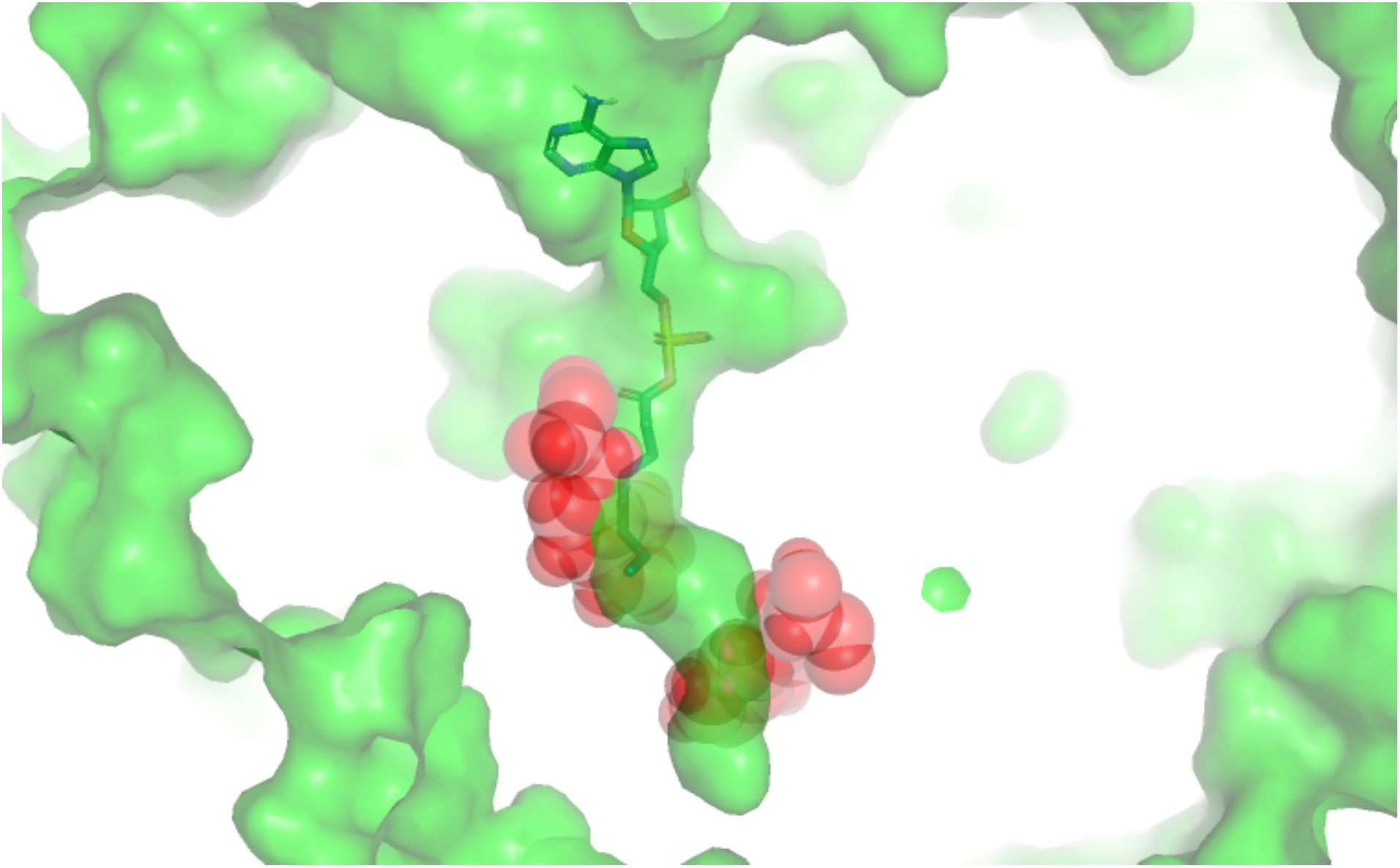
A fatty acyl-AMP ligase with substrate docked. The two red sections represent two point mutations under consideration: the location of the docked ligand shows that the upper point mutation would occlude the pictured C8 ligand, while the lower point mutation would occlude ligands C10 or larger.

Foldy was used to evaluate AlphaFold Multimer’s ability to predict chimeric polyketide synthase production. Nava et al (in review) evaluated whether a chimeric PKS’s predicted structure is indicative of its production titer. A total of 144 interacting KS-AT / ACP pairs from a seminal paper about module swaps [12] were co-folded with AlphaFold Multimer in Foldy, and the protein contact probability map was used as a proxy for likelihood of protein interaction, as described by Humphrey’s et al[6]. The authors found the predicted structures informative and worth more investigation.

Metabolic engineers used Foldy to predict the function of dozens of domains of unknown function (DUFs) in *P. putida*, including DUF1302 and DUF1329. These two DUFs which were previously suspected of being involved in hydrolase activity [13] may actually be involved in substrate transport. DUFs with pfam IDs DUF1302 and DUF1329, represented in *P. putida* by PP_0765 and PP_0766, have no predicted function, but prior RB-TnSeq experiments show their function correlates with both a periplasmic protein (PP_2018) and a multidrug efflux transporter (PP_2019) [13]. Combinatorial protein-protein docking sims, done with AlphaFold in Foldy, predicted two complexes. First, the two DUFs seem to interact with one another: PP_0765, which appears to be a membrane bound beta barrel, is predicted to form a complex with PP_0766 (Fig 5, top). Second, PP_0766 is predicted to form a heterotrimer with both PP_2018 and PP_2019 (Fig 5, bottom). This may indicate that these proteins are not in fact involved in hydrolase activity as previously reported[13] but rather are components of a novel transport system. Future work should interrogate this system in greater detail to determine the directionality of transport and *in vivo* function

**Fig 5:**
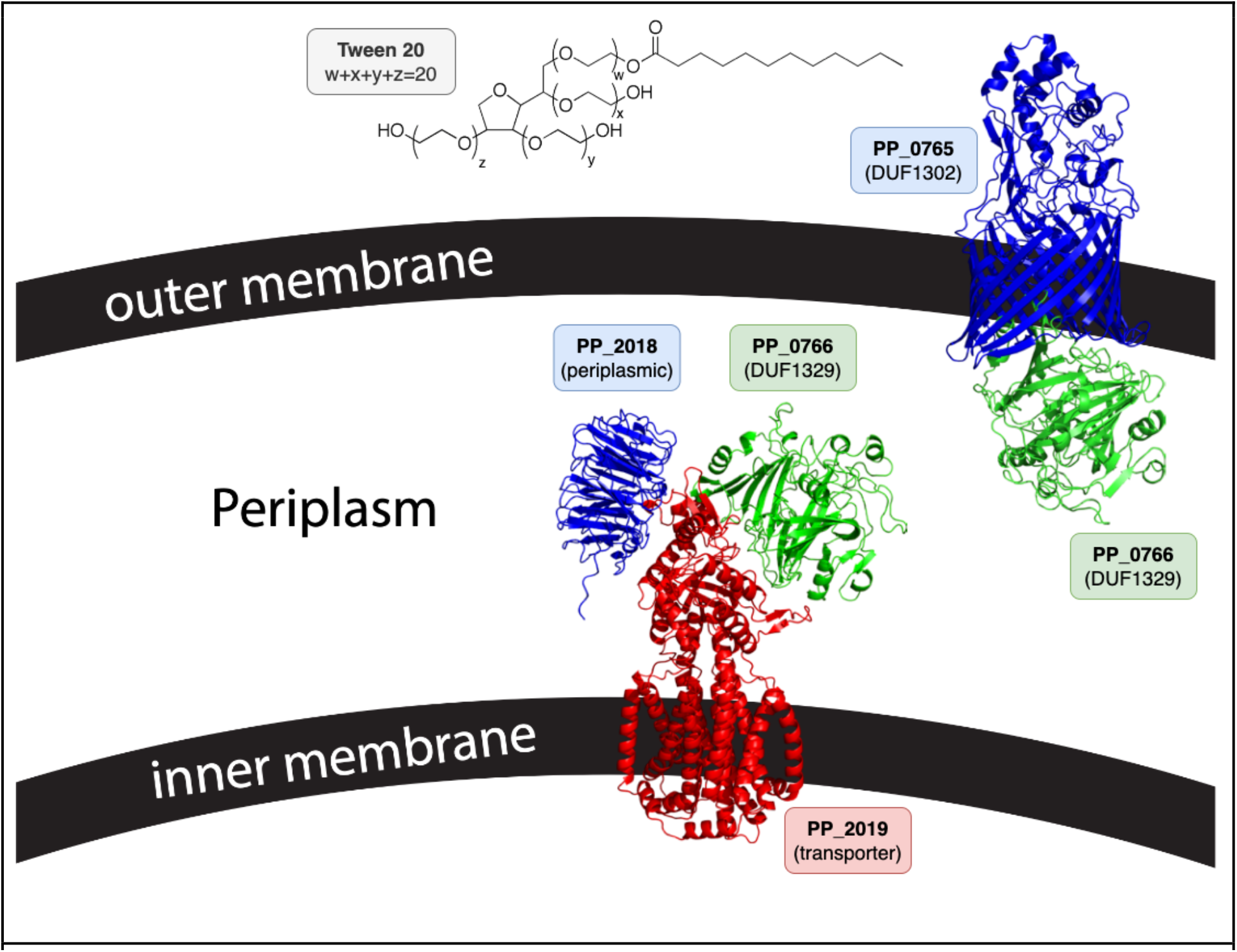
Putative complex formation of two domains of unknown function in *P. putida*. Two DUFs in *P. putida*, previously hypothesized to have hydrolase activity may actually be involved in substrate transport: PP_0765 (DUF1302, top blue) and PP_0766 (DUF1329, top green, bottom green). PP_0765 has the characteristic beta-barrel of a membrane protein, and shows high likelihood of forming a complex with PP_0766 (top). Additionally, PP_0766 is predicted to form a heterotrimer with PP_2018 (bottom blue) and PP_2019 (bottom red).

### Conclusion

Foldy is easier to use than other AlphaFold implementations, and addresses some of their limitations [1,7]. User experience studies indicate that small improvements in the enjoyability of a tool may have significant effects on tool use [14]. Foldy will increase biologists’ productivity by increasing the adoption of computational structure tools. The adoption of Foldy by institutions, including those without large compute clusters or GPUs, will make high-accuracy protein structure prediction more accessible.

## Availability and Future Developments

Website built on Kubernetes, with the Helm chart and Dockerfiles freely available at https://github.com/JBEI/foldy. One can access the “public” structures in our group’s deployment at foldy.lbl.gov.

## Acknowledgements

This work was part of the DOE Joint BioEnergy Institute (jbei.org) supported by the U.S. Department of Energy, Office of Science, Office of Biological and Environmental Research, through contract DE-AC02-05CH11231 between Lawrence Berkeley National Laboratory and the U.S. Department of Energy and by a Department of Energy Office of Science Distinguished Scientist Award to J.D.K. J.B.R. was supported in part by a fellowship award under contract [FA9550-21-F-0003] through the National Defense Science and Engineering Graduate (NDSEG) Fellowship Program, sponsored by the Air Force Research Laboratory (AFRL), the Office of Naval Research (ONR) and the Army Research Office (ARO). A.A.N. was supported by a National Science Foundation Graduate Research Fellowship, fellow ID [2018253421]. The views and opinions of the authors expressed herein do not necessarily state or reflect those of the United States Government or any agency thereof. Neither the United States Government nor any agency thereof, nor any of their employees,makes any warranty, expressed or implied, or assumes any legal liability or responsibility for the accuracy, completeness, or usefulness of any information, apparatus, product, or process disclosed, or represents that its use would not infringe privately owned rights.

J.D.K. has financial interests in Amyris, Ansa Biotechnologies, Apertor Pharma, Berkeley Yeast, Demetrix, Lygos, Napigen, ResVita Bio, and Zero Acre Farms.

